# The pH-responsive SmrR-SmrT system modulates *C. difficile* antimicrobial resistance, spore formation, and toxin production

**DOI:** 10.1101/2023.11.02.565354

**Authors:** Daniela Wetzel, Zavier A. Carter, Marcos P. Monteiro, Adrianne N. Edwards, Shonna M. McBride

## Abstract

*Clostridioides difficile* is an anaerobic gastrointestinal pathogen that spreads through the environment as dormant spores. To survive, replicate, and sporulate in the host intestine, *C. difficile* must adapt to a variety of conditions in its environment, including changes in pH, the availability of metabolites, host immune factors, and a diverse array of other species. Prior studies showed that changes in intestinal conditions, such as pH, can affect *C. difficile* toxin production, spore formation, and cell survival. However, little is understood about the specific genes and pathways that facilitate environmental adaptation and lead to changes in *C. difficile* cell outcomes. In this study, we investigated two genes, *CD2505* and *CD2506,* that are differentially regulated by pH to determine if they impact *C. difficile* growth and sporulation. Using deletion mutants, we examined the effects of both genes (herein *smrR* and *smrT*) on sporulation frequency, toxin production, and antimicrobial resistance. We determined that SmrR is a repressor of *smrRT* that responds to pH and suppresses sporulation and toxin production through regulation of the SmrT transporter. Further, we showed that SmrT confers resistance to erythromycin and lincomycin, establishing a connection between the regulation of sporulation and antimicrobial resistance.

**IMPORTANCE:** *C. difficile* is a mammalian pathogen that colonizes the large intestine and produces toxins that lead to severe diarrheal disease. *C. difficile* is a major threat to public health due to its intrinsic resistance to antimicrobials and its ability to form dormant spores that are easily spread from host to host. In this study, we examined the contribution of two genes, *smrR* and *smrT* on sporulation, toxin production, and antimicrobial resistance. Our results indicate that SmrR represses *smrT* expression, while production of SmrT increases spore and toxin production, as well as resistance to antibiotics.

## INTRODUCTION

*C. difficile* is a leading cause of antibiotic-associated diarrhea throughout the world (1, 2). Infections by *C. difficile* are easily transmitted due to the ability of the bacterium to form spores during transit through the host intestine. The spore form provides *C. difficile* extraordinary protection from the effects of oxygen, most disinfectants, and antibiotics (3–5). However, the conditions within the host and the bacterial factors that activate *C. difficile* spore formation are poorly understood.

One known environmental trigger of *C. difficile* sporulation is pH. *C. difficile* resides and replicates within the large intestine, where the pH can range from 5.2 to 7.8 (6). The pH of the colon can also vary with the health and diet of the host (6, 7). *C. difficile* can adapt to the changing pH of the intestine and can even modulate host pH (8, 9). Conditions of both relatively high and low physiological pH can prompt sporulation in different *C. difficile* strains (10). But, it is less clear which factors encoded by *C. difficile* respond to changes in pH to affect sporulation initiation.

In this study, we investigated the effects of two *C. difficile* genes that are regulated in response to changes in pH, *CD2505* and *CD2506,* which encode a transcriptional regulator and an associated transporter. We tested the hypothesis that these genes are involved in pH-dependent sporulation responses and examined the functions of both factors. We determined that CD2505 regulates the expression of both genes and suppresses sporulation by repressing expression of the *CD2506* transporter. In addition, we found that the CD2506 transporter confers resistance to antibiotics, thereby linking antibiotic resistance and spore formation. Based on these results, we renamed the genes SmrR and SmrT for their respective roles as a repressor and transporter that modulate sporulation and macrolide/lincosamide resistance.

## RESULTS

### Expression of the *smr* operon is regulated by pH

In a prior investigation, we found that *C. difficile* sporulation frequency increased with corresponding increases in the environmental pH, which accompanied global changes in transcription (10). One locus that changed considerably with pH variation was a predicted two-gene operon, *CD2505-CD2506 (smrR-smrT)*, which demonstrated a significant decrease in expression as pH increased (**Figure 1**). SmrR is annotated as a putative AcrR-like transcriptional regulator of the TetR-family, while SmrT is predicted to encode an LmrB-like transporter of the Major Facilitator Superfamily (MFS). AcrR-like regulators are transcriptional repressors that respond to a variety of toxic substances and ions, and often repress the expression of antimicrobial transporters (11–14). The LmrB-subfamily of transporters are drug/H^+^ antiporters that can confer resistance to a wide range of toxic substances (15–18).

**Figure 1.**
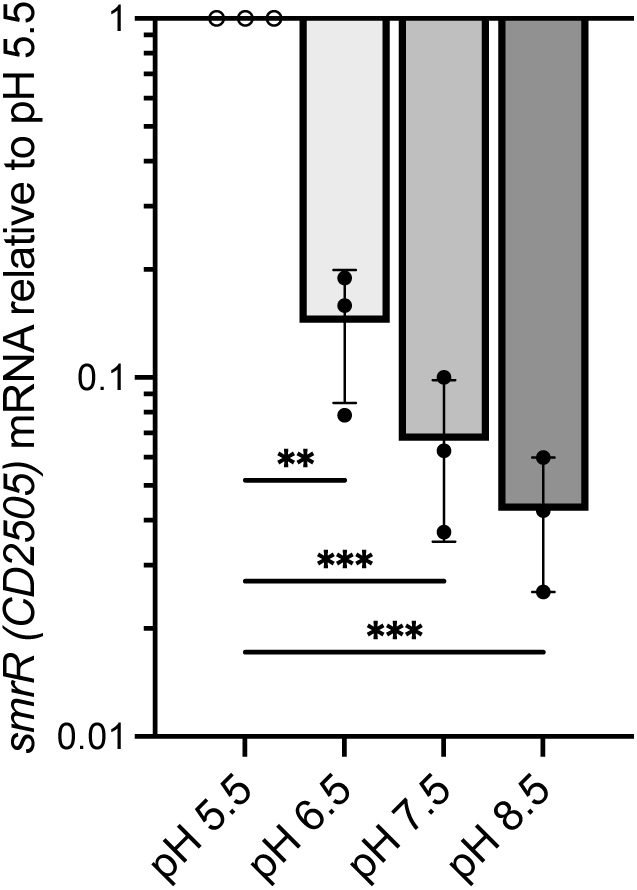
Expression of *smrR* is pH-dependent. qRT-PCR analysis of *smrR* (*CD2505;* TetR-like regulator) expression in 630Δ*erm* grown on 70:30 sporulation agar at pH 5.5, 6.5, 7.5, and 8.5 and sampled at H12, respectively. Expression levels were analyzed by one-way ANOVA and Dunnett’s multiple comparisons test compared to the pH 5.5. \*\**P* ≤ 0.01, \*\*\**P* ≤ 0.001

Based on the proximity and arrangement of *smrR* and *smrT*, we predicted that these genes constitute a dicistronic operon (**Figure S1**). To examine co-transcription of *smrR-smrT*, we used cDNA generated from *C. difficile* strain 630Δ*erm* grown on sporulation medium and tested for transcriptional linkage by PCR. A product was amplified across the *smrR-smrT* open reading frames, indicating that these genes are co-transcribed (**Figure S1**).

### SmrR suppresses sporulation of *C. difficile*

As both spore formation and *smrRT* expression correlate with changes in pH, we hypothesized that SmrR and/or SmrT may affect sporulation in *C. difficile.* To test this, we created deletion mutants in *smrR* and *smrT* by allelic exchange (19) (**Figure S2**). The resulting mutants were then examined for the ability to produce spores in a modified 70:30 sporulation broth (pH 7.2) without Tris, to reduce buffering capacity (20–22). As shown in **Figure 2**, the Δ*smrR* regulator mutant (MC1681) exhibited significantly greater sporulation frequency than the parent strain (WT, 630Δ*erm*), while the Δ*smrT* transporter mutant (MC1682) had no change in sporulation. The Δ*smrR* phenotype was fully complemented when *smrRT* was reintroduced and expressed on an extrachromosomal plasmid, implicating SmrR as a suppressor of sporulation (**Figure S3**).

**Figure 2.**
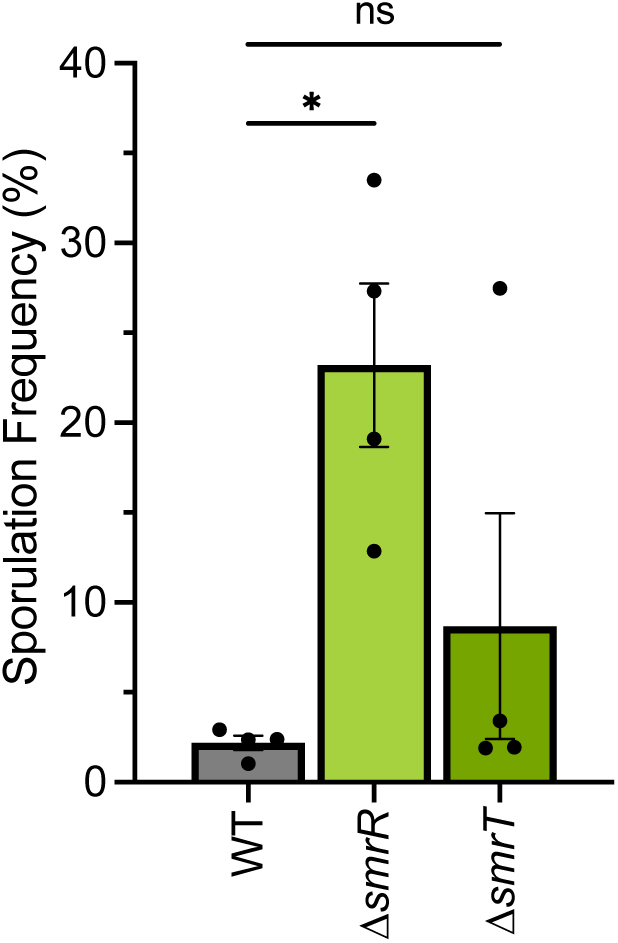
The *smrR* mutant has increased spore formation. Sporulation frequency of wild-type (630Δ*erm*), the *CD2505* mutant (MC1681), and the *CD2506* mutant (MC1682) grown in 70:30 broth (pH 7.2) for 24 h. The means, individual data points, and standard deviations are shown for four independent replicates. Data were analyzed by one-way ANOVA and Dunnett’s multiple comparison test comparing the mutant strains to 630Δ*erm*. \**P* ≤ 0.05; ns: not significant.

To determine if the increase in sporulation observed for the 630Δ*erm smrR* mutant extends to other *C. difficile* isolates, we examined the impact of *smrR* on sporulation in the epidemic UK1 strain (027 ribotype). Using a CRISPR interference (CRISPRi) approach (23, 24), we knocked-down transcription of the *smrR* ortholog in UK1 (*CDR20291_2397*) As illustrated in **Figure S4**, when *smrR* transcript level was decreased by CRISPRi, a dramatic increase in sporulation frequency was observed relative to the control strain, indicating that SmrR comparably impacts sporulation in divergent *C. difficile* strains.

### SmrR exclusively regulates the *smrRT* operon

Since SmrR encodes an apparent transcriptional regulator, we considered that it likely regulated the transcription of another factor(s) to effect changes in sporulation. To determine the SmrR regulon, we assessed transcription during growth in sporulation broth by RNA-seq for the *smrR* null mutant, relative to the parent strain (**Table 1**). We observed differential expression of one transcript in the *smrR* mutant—*smrT—*which was more than 6-fold greater in the *smrR* mutant. Given that no additional transcripts were differentially expressed in the *smrR* mutant, we concluded that SmrR represses only *smrR-smrT* transcription under the conditions tested.

**Table 1.**
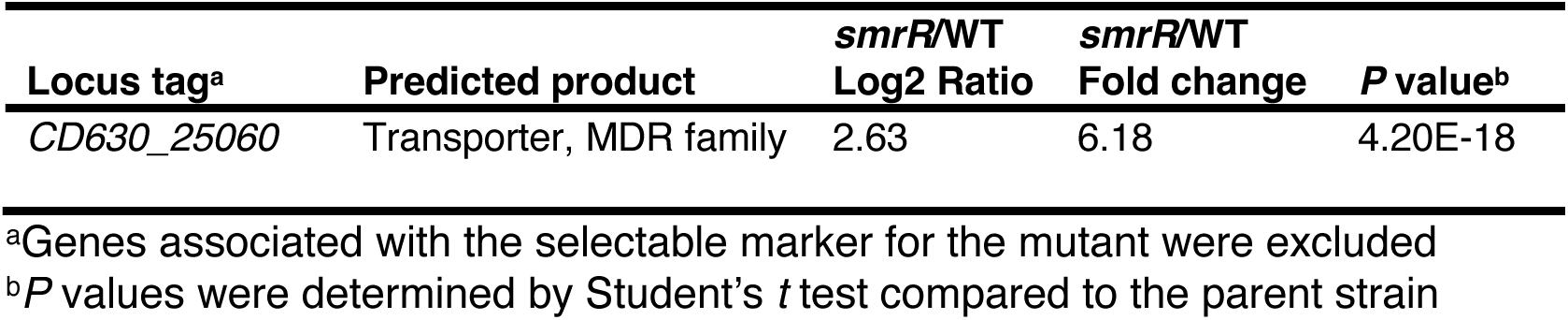
Genes differentially expressed in the Δ*smrR* mutant by RNA-seq.

To further define regulation of the *smrRT* operon, a transcriptional reporter fusion of the *smr* promoter and the alkaline phosphatase gene was constructed (P*smr::phoZ*) and expression assessed in the wild-type and *ΔsmrR* strains grown in unbuffered sporulation medium at pH 6.2 (**Figure 3**). Reporter expression from P*smr* was pronounced in the parent strain carrying the fusion, in agreement with the transcription results for *smrR* under acidic conditions (**Figure 1**). However, reporter expression was markedly higher in the Δ*smrR* mutant, which exhibited 8.9-fold greater activity than the parent strain. The difference in reporter expression from P*smr* for the *smrR* mutant and parent strain (**Figure 3**) was similar to the increased *smrT* expression that was observed for the Δ*smrR* mutant by RNA-seq (**Table 1**). P*smr::phoZ* activity was unchanged by growth stage in the parent strain, suggesting that transcriptional repression by SmrR is not subject to regulation by stationary phase conditions or factors (**Figure S5**). Together these data are consistent with SmrR acting as the primary transcriptional regulator that represses expression of the *smrRT* operon.

**Figure 3.**
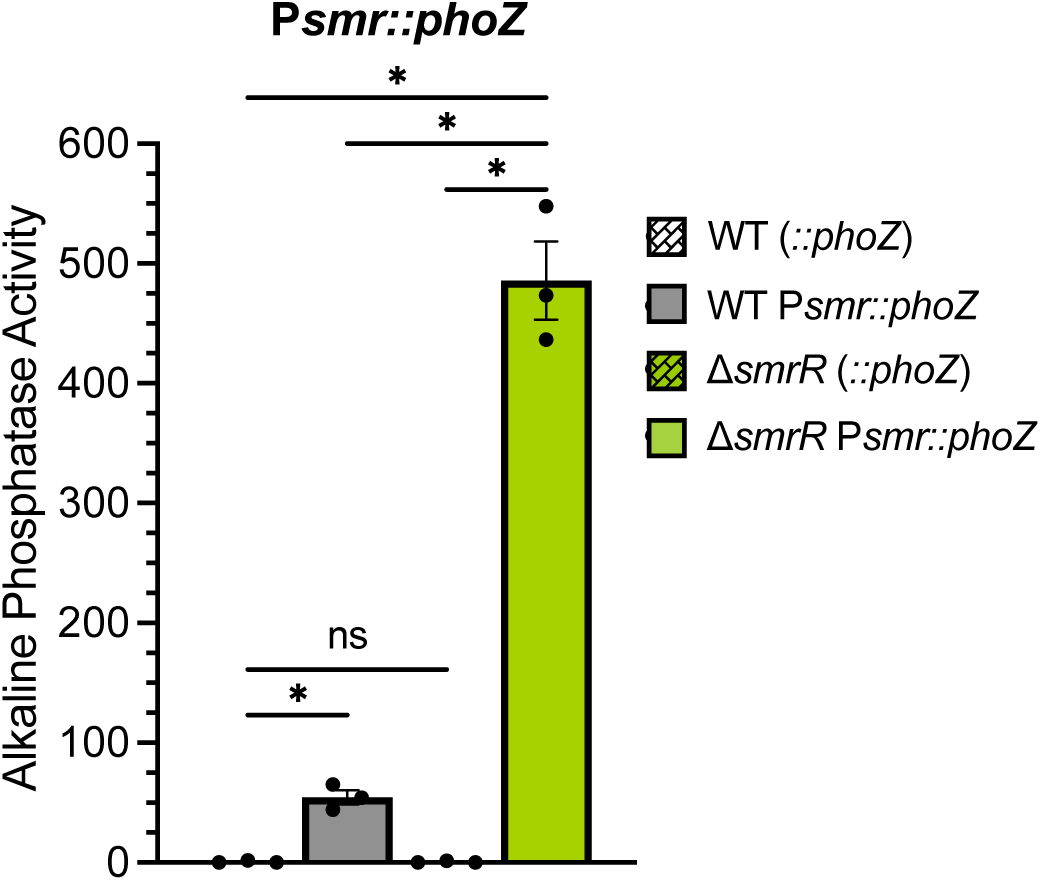
SmrR represses expression from the *smr* promoter. Alkaline phosphatase activity of the P*smr::phoZ* fusion in *C. difficile* strain 630Δ*erm* (WT, MC1775), the *smrR* mutant (Δ*smrR,* MC1777), and respective promoterless control strains (WT, MC448 and Δ*smrR*, MC1776). Strains were grown in 70:30 broth, pH 6.2 with 1 *µ*g/ml thiamphenicol and samples were taken during logarithmic growth (OD_600_ 0.5). The means, individual data points, and standard deviations are shown for three independent replicates. Data were analyzed by two-way ANOVA with Tukey’s multiple comparison test. \**P* ≤ 0.05; ns: not significant.

### Over-expression of the SmrT transporter increases spore formation and toxin production

Because *smrT* was the only transcript differentially regulated in the Δ*smrR* mutant, we considered that the basis for increased sporulation of the Δ*smrR* strain was over-expression of the transporter, SmrT. To test this hypothesis, we generated a construct to express *smrT* under a nisin-inducible promoter in the wild-type background (MC2508; 630Δ*erm,* P*cpr::smrT*). Sporulation frequency was then assessed in the *smrT* over-expressing strain and vector control (**Figure 4**). Over-expression of *smrT* resulted in ∼7.3-fold increase in spore formation relative to the control. These results suggest that increased transport of substrates and/or ions via SmrT triggers *C. difficile* to sporulate.

**Figure 4.**
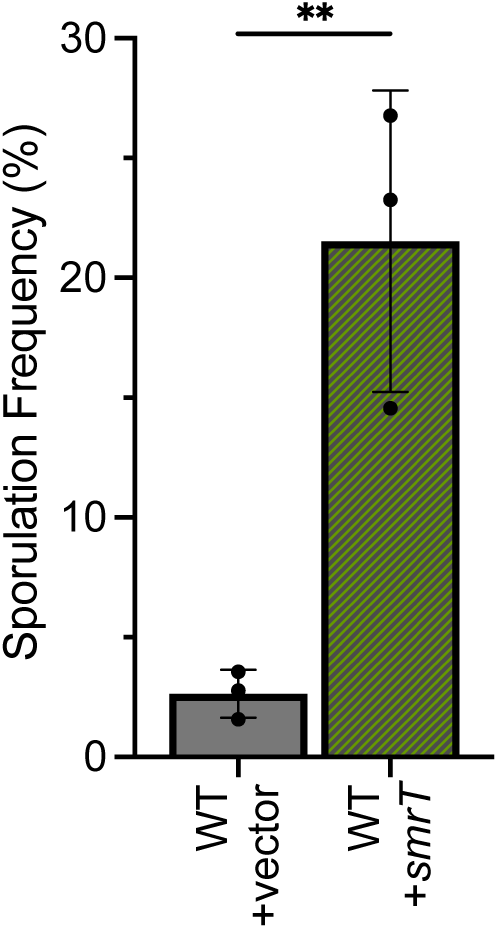
Overexpression of *smrT* increases spore formation. Sporulation frequency of the wild-type control with vector (MC282; 630Δ*erm*, pMC211) and overexpressing *smrT* (MC2508; 630Δ*erm*, pMC1282). Strains were grown in 70:30 broth, pH 7.2 with 1 *µ*g/ml thiamphenicol and 0.1 *µ*g/ml nisin and assessed for spore formation by phase contrast microscopy. The means, individual data points, and standard deviations are shown for three independent replicates. Data were analyzed by paired Student’s *t* test. \**P* ≤ 0.05.

The initiation of sporulation and the production of the toxins TcdA and TcdB are often concurrent due to overlapping regulation (25, 26). Although no difference in toxin expression was observed in the Δ*smrR* mutant during exponential growth in sporulation medium (**Table 1**), toxin production was significantly increased in the supernatant following 24 h of growth in the standard toxin quantification medium, TY (**Figure 5**). However, transcription of sporulation and toxin genes was not observed in the Δ*smrR* mutant during active growth in sporulation medium (**Table 1**), suggesting that a greater fraction of the population sporulates in response to elevated SmrT, but sporulation and toxin production do not occur prematurely. These data suggest that however increased SmrT perturbs the cell to increase toxin production, it follows shared regulation that leads to increased sporulation.

**Figure 5.**
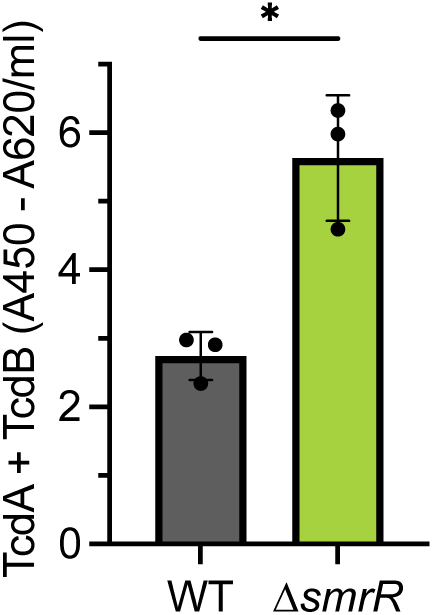
The *smrR* mutant has increased toxin production. Quantification of TcdA and TcdB from supernatants of 630Δ*erm* (WT) and the *smrR* mutant (MC1681) grown in TY for 24 h. The means, individual data points, and standard deviations are shown for three independent replicates. Data were analyzed by paired Student’s *t* test. \**P* ≤ 0.05.

### SmrT confers resistance to macrolide and lincosamide antibiotics

SmrT is an LmrB-like drug/H^+^ antiporter, which are known to confer resistance to a variety of antimicrobial compounds. SmrT shares 61% similarity / 38% identity with LmrB of *Bacillus subtilis* (BLASTp); however, SmrT has not been implicated in *C. difficile* resistance to antimicrobials and its function is unverified. We examined resistance of the *smrR* and *smrT* mutants to several antimicrobials that are associated with similar transporters, including lincomycin, erythromycin, spectinomycin, and ethidium bromide (18, 27–29). The minimal inhibitory concentration (MIC) of the Δ*smrR* mutant for lincomycin (a lincosamide) was more than 20-fold greater, while the MIC for erythromycin (a macrolide) was more than 645-fold more than the parent strain (**Table 2**). Surprisingly, the inverse was not true for the Δ*smrT* mutant, which exhibited no apparent change in MIC for any of the antimicrobials tested. No change in resistance to spectinomycin or ethidium bromide were observed for either mutant. Thus, the absence of SmrT had negligible impact on antimicrobial resistance, but over-expression of the transporter dramatically increased *C. difficile* resistance to both lincomycin and erythromycin. These results are similar to the effects observed for sporulation with *smrT* deletion and over-expression.

**Table 2.** MIC values for Δ*smrR* and Δ*smrT* mutants.

| Antimicrobial | WT | $\Delta smrR$ | $\Delta smrT$ |
| --- | --- | --- | --- |
| Lincomycin | 25-50 | 1000 | 25-50 |
| Erythromycin | 3.1 | >2000 | 3.1 |
| Spectinomycin | 1000 | 1000 | 1000 |
| Ethidium Bromide | 25 | 25 | 25 |

The increase in antimicrobial resistance for the Δ*smrR* mutant suggested that SmrR may respond to antimicrobials directly to regulate expression of the *smrRT* operon. To this end, we further investigated the transcription from the *smr* promoter in an assortment of antimicrobials using the P*smr::phoZ* reporter (**Figure S6**). While the wild-type showed modest increases in activity in the presence of erythromycin and lincomycin, the differences were not significant when adjusted for multiple comparisons. No differences in activity were observed with any substance for the Δ*smrR* mutant, again supporting SmrR as the primary regulator of *smrRT* expression.

## DISCUSSION

The primary objective of this work was to determine if the *smrRT* system functions as a link between the response of *C. difficile* to pH variation and sporulation. However, the results we obtained indicate that SmrRT not only connects sporulation to environmental pH, but it also links antimicrobial resistance with these physiological responses. We found that increased levels of SmrT promotes sporulation, while SmrR reduces sporulation through repression of *smrRT* expression. Despite the increase in sporulation when SmrT was over-expressed, sporulation was not significantly impacted by removal of *smrT*, indicating that SmrT can promote sporulation, but is not needed for sporulation to occur. Further, growth and adaptation to pH (not shown) was not affected in either the Δ*smrR* or the Δ*smrT* mutant, suggesting that the *smr* locus is not involved in altering the extracellular pH.

*C. difficile* is well known for resistance to a wide variety of therapeutic antimicrobials including tetracyclines, ß-lactams, aminoglycosides, macrolides, lincosamides, and fluoroquinolones (30–34). Despite the considerable impact of antimicrobial resistance on the spread of CDI, *C. difficile* antibiotic resistance profiles are rarely tested in clinical settings and the resistance mechanisms to non-CDI antimicrobials are understudied. While *C. difficile* strains are often associated with resistance to macrolides and lincosamides, the only resistance determinants known to confer significant resistance to these antibiotics in *C. difficile* are the *erm* (MLS) family elements that act via ribosomal methylation (35–37). Prior to the development of *C. difficile* genetic manipulation techniques, the predicted *C. difficile* efflux gene, *cme,* was examined for resistance functions in heterologous hosts, but resistance to erythromycin by Cme was negligible (38). The same study attempted to determine if SmrT (annotated as LinCD and *lind*) provided resistance to macrolides; however, they were unable to test its function due to lack of expression of the gene in *E. coli, Enterococcus faecalis,* or *Staphylococcus aureus* (38). Resistance to macrolides by efflux has been proposed for strains that do not encode apparent *erm* mechanisms, but the genetic elements involved were not identified (39). Our data suggest that the robust resistance to macrolides and lincosamides observed in the Δ*smrR* mutant could account for high-level resistance observed in *C. difficile* isolates that lack *erm*/MLS resistance genes. Macrolide and lincosamide treatments may select for mutations in *smrR* or the SmrR-binding region of P*smr*, which would relieve repression of *smrRT,* since increased expression of SmrT would confer significant resistance.

Several questions remain about SmrRT, including how it effects sporulation in the host and how the intestinal environment affects *smrRT* regulation. We observed robust expression of *smrR* in low pH conditions, suggesting that low intestinal pH could trigger SmrR derepression of the *smr* promoter. Since expression from the *smr* promoter was not significantly increased by either erythromycin or lincomycin, these antibiotics probably do not impact the typical regulation of this operon in the intestine. In addition to regulation by pH, data from other studies suggest that *smrRT* expression may be influenced by nutritional availability. Expression of *smrR-smrT* was found to increase 4-6 fold in the presence of glucose and was partially repressed by the carbon catabolite regulator, CcpA (40). In addition, expression of *smrRT* decreased about 5-fold in a *codY* mutant, suggesting that the availability of branched-chain amino acids or GTP may promote transcription and activity of the transporter (41). CcpA and CodY each repress toxin and sporulation, which are both impacted by SmrT, but how SmrR-SmrT would impact regulation through these regulators is not apparent. Further study is needed to understand function of SmrR and SmrT in the host, as well as their roles in pathogenesis.

## MATERIALS AND METHODS

### Bacterial strains and growth conditions

Bacterial strains and plasmids used in this study are listed in **Table 3**. An anaerobic chamber (Coy Laboratory Products) was used to cultivate *C. difficile* in an atmosphere of 10% H_2_, 5% CO_2_, and 85% N_2_, at 37°C (42, 43). Strains were routinely grown fresh from stock in brain heart infusion-supplemented with 0.5% yeast extract (Βecton Dickinson Company) broth or agar plates. Cultures were supplemented with 0.1% taurocholate to induce germination and 0.2% fructose to prevent sporulation, whenever needed. Strains were grown in the presence of 1-5 *µ*g/ml thiamphenicol, 2-5 *µ*g/ml erythromycin, or 100 ng/ml anhydrotetracycline (ATc), when needed for plasmid maintenance or selection (Sigma-Aldrich). *Escherichia coli* was grown at 37°C in LB medium with 100 μg/ml ampicillin and/or 20 μg/ml chloramphenicol (Sigma-Aldrich), as needed for plasmid maintenance. Growth, and pH were measured using a spectrophotometer and a portable pH/ORP meter (HI98190, HANNA instruments), respectively.

**Table 3.** Bacterial Strains and plasmids.

| Plasmid or Strain | Relevant genotype or features | Source, construction or reference |
| --- | --- | --- |
| **Strains** |  |  |
| *E. coli* |  |  |
| DH5α Max Efficiency | F<sup>-</sup> Φ80*lacZ*ΔM15 Δ(*lacZYA-argF*)<br>U169 *recA1 endA1 hsdR17* (rk<sup>-</sup>, mk<sup>+</sup>) *phoA supE44 λ*<sup>-</sup><br>*thi1 gyrA96 relA1* | Invitrogen |
| HB101 pRK24 | F<sup>-</sup> *mcrB mrr hsdS20*(r<sub>B</sub><sup>-</sup> m<sub>B</sub><sup>-</sup>) *recA13 leuB6 ara-14 proA2 lacY1 galK2 xyl-5 mtl-1 rpsL20* | B. Dupuy |
| *C. difficile* |  |  |
| 630Δ*erm* | Erm<sup>S</sup> derivative of strain 630 (ribotype 012) | Nigel Minton, (56) |
| UK1 | Clinical isolate (ribotype 027) | (57) |
| MC282 | 630Δ*erm* pMC211 | (21) |
| MC310 | 630Δ*erm* *spo0A::erm* | (21) |
| MC324 | 630Δ*erm* pMC123 | (21) |
| MC448 | 630Δ*erm* pMC358 | (53) |
| MC855 | 630Δ*erm* *spo0A::erm* pMC123 | (58) |
| MC1681 | 630Δ*erm* Δ*smrR::ermB* | This study |
| MC1682 | 630Δ*erm* Δ*smrT* | This study |
| MC1771 | MC1681 Δ*smrR* pMC123 | This study |
| MC1775 | 630Δ*erm* pMC1019 | This study |
| MC1776 | 630Δ*erm* Δ*smrR* pMC358 | This study |
| MC1777 | 630Δ*erm* Δ*smrR* pMC1019 | This study |
| MC1861 | MC1681 Δ*smrR* pMC1059 | This study |
| MC2186 | UK1 pMC1123 | This study |
| MC2263 | UK1 pMC1178 | This study |
| MC2508 | 630Δ*erm* pMC1282 | This study |
| **Plasmids** |  |  |
| pIA33 | P<sub>*xyl*</sub>::*dCas9-opt* P<sub>*gdh*</sub>::*sgRNA-rfp catP* | (23) |
| pJIR1457 | *ermB oriCP oriEC oriT* | (59) |
| pMSR | pseudo-suicide plasmid used for allelic exchange | (19) |
| pRK24 | Tra<sup>+</sup>, Mob<sup>+</sup>; *bla*, *tet* | (60) |
| pMC123 | *E. coli*-*C. difficile* shuttle vector; *bla*, *catP* | (49) |
| pMC211 | pMC123 with *cprA* promoter, *bla catP* | (21) |
| pMC358 | pMC123::*phoZ* | (53) |
| pMC914 | pMSR with flanking homology for *CD2506* deletion | This study |
| pMC946 | pMSR with *CD2505* homology regions flanking *ermB* | This study |
| pMC1019 | pMC123 with *Psmr::phoZ* | This study |
| pMC1059 | pMC123 with *PsmrRT* | This study |
| pMC1123 | P<sub>*cprA*</sub>::*dCas9-opt* P<sub>*gdh*</sub>::*sgRNA-neg catP* | (24) |
| pMC1178 | P<sub>*cprA*</sub>::*dCas9-opt* P<sub>*gdh*</sub>::*sgRNA-CDR20291_2397 catP* | This study |
| pMC1282 | pMC211 with *Pcpr::smrT* | This study |

### Strain and plasmid construction

Primer design and the template for PCR reactions were based on *C. difficile* strain 630, unless otherwise noted (GenBank accession NC_009089.1). Genomic DNA (gDNA) was isolated from *C. difficile* strains using a modified Bust ‘N’ Grab protocol (44, 45). The Benchling CRISPR Guide RNA Design tool was used to create the sgRNA targeting *CDR20291_2397*, which was generated by PCR (23, 24). The details of vector construction are provided in **Supplemental Table S1**. The oligonucleotide primers used in this study are listed in **Table 4**. Plasmids were introduced to *C. difficile* by conjugation with *E. coli*, as previously described (46). After conjugation with *C. difficile*, *E. coli* was counterselected using 100 *µ*g/ml kanamycin and gene deletions selected and screened for as previously described (19, 47).

**Table 4.**
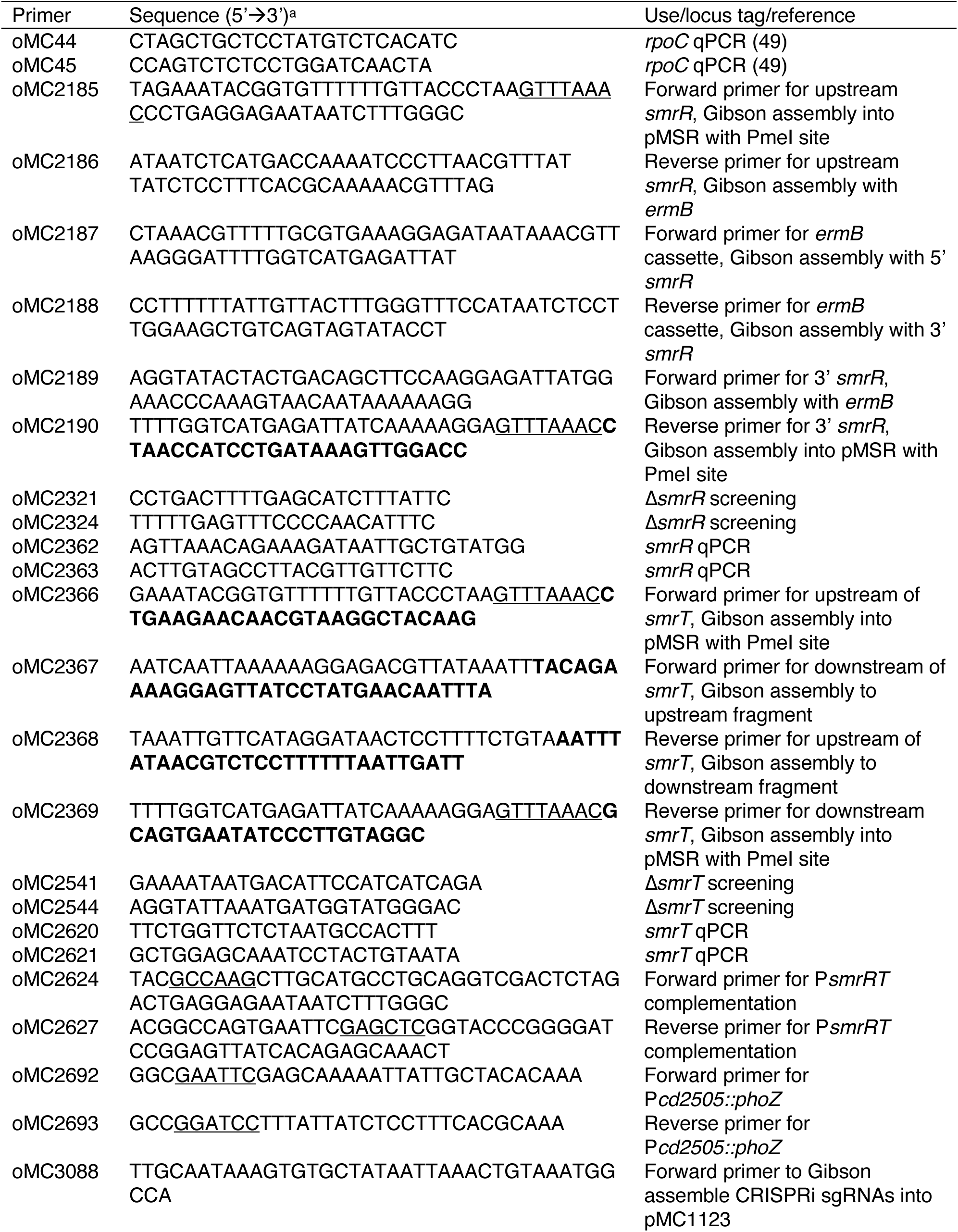
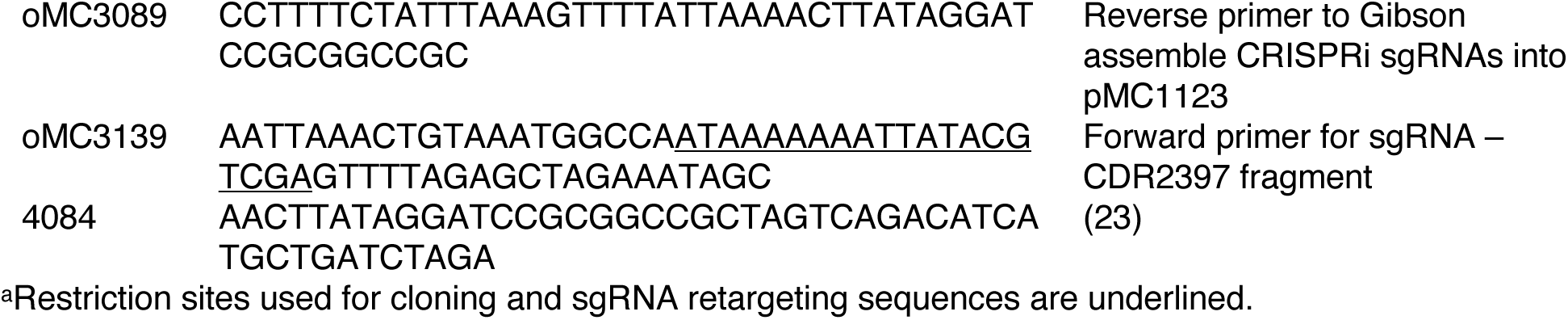
Oligonucleotides.

### Quantitative reverse transcription PCR (qRT-PCR)

*C. difficile* cultures were grown on 70:30 sporulation agar (-Tris) at pH 5.5, 6.5, 7.5, and 8.5, respectively, harvested after 12 h (H12) into ice-cold water:ethanol:acetone (3:1.5:1.5), and stored at −70°C. RNA isolation, DNase I treatment, and cDNA synthesis was performed as previously explained (21, 48, 49). Quantitative reverse-transcription PCR (qRT-PCR) was performed in technical triplicate with 50 ng of cDNA per reaction using the SensiFAST SYBR & Fluorescein Kit from Bioline, on a Roche Lightcycler 96. Samples containing RNA without the addition of reverse transcriptase were used to control for contaminating genomic DNA. Assay results (Ct values) were averaged and analyzed by the comparative cycle threshold method with the housekeeping *rpoC* transcript as a normalizer (50). The data are presented as the means, individual values, and standard deviation of the means for at least three independent biological replicates. For expression by pH condition, a one-way ANOVA and Dunnett’s test was performed for statistical comparison to pH 5.5 using GraphPad Prism version 10.0.2.

### Sporulation assays

Sporulation assays were performed as previously described in 70:30 sporulation medium, without the addition of Tris base and pH adjusted as noted (22, 51). UK1 sporulation assays were performed on standard 70:30 agar supplemented with 2 *µ*g/ml thiamphenicol and 1 *µ*g nisin for plasmid maintenance and induction of expression, respectively. Sporulation frequency was determined by ethanol resistance and/or phase-contrast microscopy (22, 51). A *spo0A* mutant (strain MC310, sporulation defective) was used as a negative sporulation control in all assays. Statistical significance was determined by one-way ANOVA with Dunnett’s or Sidak’s post-test, or Student’s *t* test, as indicated for the specified comparison using GraphPad Prism version 10.0.2.

### RNA sequencing analysis (RNA-seq)

*C. difficile* strain 630Δ*erm* and the *smrR* mutant (MC1681) were grown in 70:30 sporulation broth (-Tris, pH 7.2) and harvested during logarithmic growth for RNA, as described for qRT-PCR. Following DNase I treatment, samples were submitted to the Microbial Genomics Sequencing Center (MiGS) for library preparation using the Illumina Stranded Total RNA prep Ligation with Ribo-Zero Plus kit and 10 bp IDT for Illumina indices. Sequencing of the RNA was performed using a NextSeq2000 instrument to yield 2x50 bp reads. Demultiplexing, quality control, and adapter trimming was performed with bcl-convert (v3.9.3; Illumina). Geneious Prime v2022.2.2 was used to map the reads to the reference genome (630; NC_009089.1). The expression levels were calculated and then subsequently compared using DESeq2 (52). DESeq2 utilizes the Wald test to calculate *P* values which are then adjusted using the Benjamini-Hochberg test (52). RNA-seq raw sequence read files were deposited to the NCBI Sequence Read Archive (SRA) BioProject PRJNA1017478.

### Alkaline phosphatase reporter assays

*C. difficile* strains containing the alkaline phosphatase (AP) transcriptional reporter fusions were grown in 70:30 medium (-Tris) at pH 6.2 and cells were harvested during logarithmic or stationary phase, as indicated. AP assays were performed as previously described (53), without chloroform. Technical duplicates were averaged for each sample, and the results provided as the individual data, mean, and standard deviation of the mean for three biological replicates. Statistical significance was determined by either a one-way ANOVA, two-way ANOVA with Tukey’s multiple comparison’s test or Student’s *t* test, as indicated for the specified comparison, using GraphPad Prism version 10.0.2.

### Minimal Inhibitory Concentrations (MIC)

The minimal inhibitory concentrations for antimicrobials against *C. difficile* strains was performed by microdilution assay in Mueller Hinton broth (BD Difco), as previously described (54, 55). Briefly, active cultures were grown to mid-logarithmic phase in MH broth (OD_600_ of 0.45), diluted 1:10, and 15 *µ*L inoculated into pre-reduced U-bottom microtiter plates containing 135 *µ*L of antimicrobials at 2-fold dilutions in MH broth. Antimicrobials tested included lincomycin (Sigma), erythromycin (Sigma), spectinomycin (Thermo-Scientific), and ethidium bromide (Sigma). Each assay was performed in technical duplicate. Negative and positive controls for contamination and growth, respectively, were included. Assays were performed for a minimum of three biological replicates and the MIC was recorded as the lowest concentration of an antimicrobial for which no visible growth was observed.

### Detection of *C. difficile* Toxin A and B

Strains were grown in BHIS broth to an OD_600_ of 0.5 and diluted 1:10 into TY medium (pH 7.4). After 24 h, cells were harvested and the supernatant assayed using a kit for the simultaneous detection of *C. difficile* toxins A and B from TGCbiomics (catalog no. TGC-E001-1), according to the manufacturer’s instructions. Technical duplicates were averaged and normalized per ml of supernatant. The results represent three independent experiments and are presented as the individual data, the means, and the standard deviation of the means. Statistical significance was determined using a two-tailed Student’s *t*-test comparing the mutant to the parent strain using GraphPad Prism version 10.0.2.

## ACKNOWLEDGEMENTS

We give thanks to members of McBride lab for helpful suggestions and discussions during the course of this work. This research was supported by the U.S. National Institutes of Health through research grants AI116933 and AI156052 to S.M.M and AI179158 to MPM. The content of this manuscript is solely the responsibility of the authors and does not necessarily reflect the official views of the National Institutes of Health.

---

**Figure S1.**
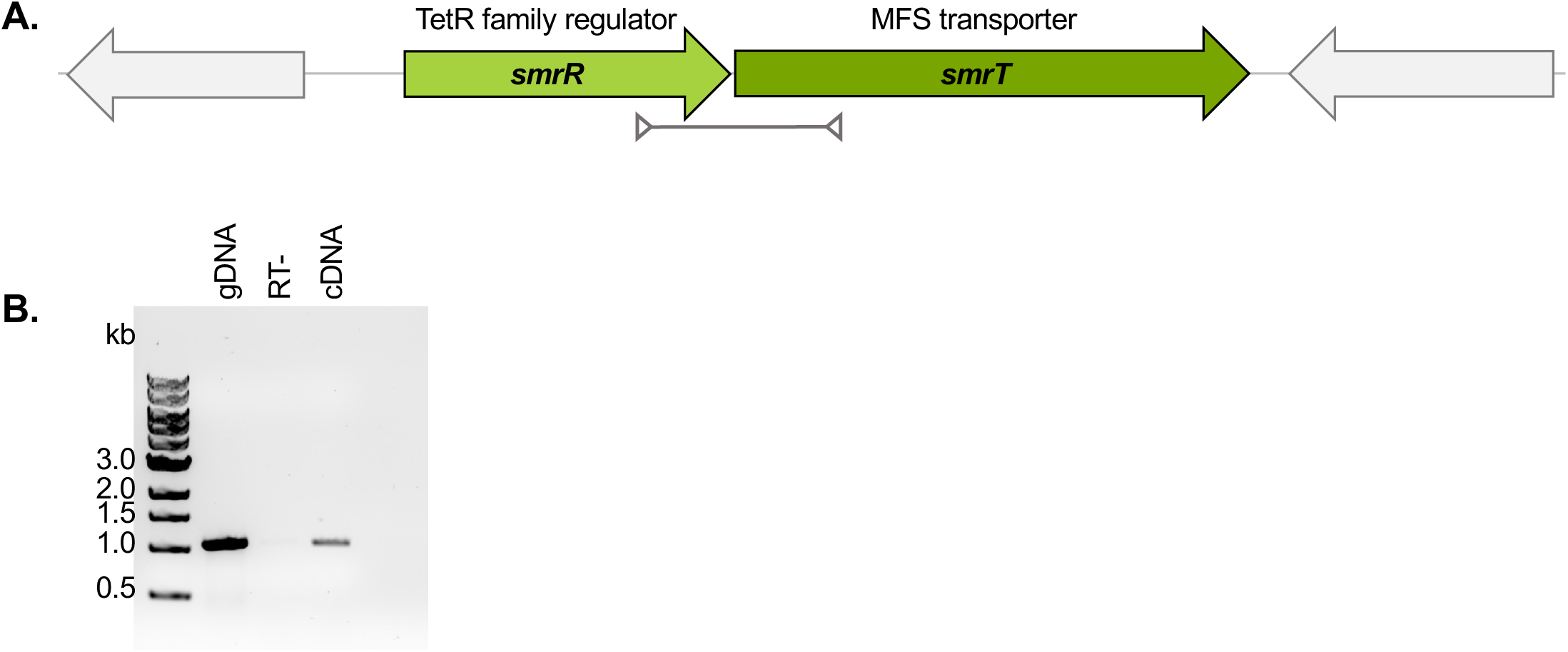
*smrR* and *smrT* are transcribed as an operon. **A)** Organization of *smrR (CD2505)* and *smrT (CD2506)* in the 630 chromosome. **B)** Transcriptional units were examined by amplification of products from adjacent open reading frames. Cultures of strain 630&*erm* were grown on sporulation agar and samples were collected after 12 h for RNA and cDNA synthesis. PCR was performed using 50 ng genomic DNA (gDNA, positive control), 50 ng cDNA templates, or 50 ng cDNA without reverse transcriptase (RT-, negative control). The 1104 bp product was generated using primers for the 39 end of *CD2505* (oMC2362) and 59 end of *CD2506* (oMC2621).

**Figure S2.**
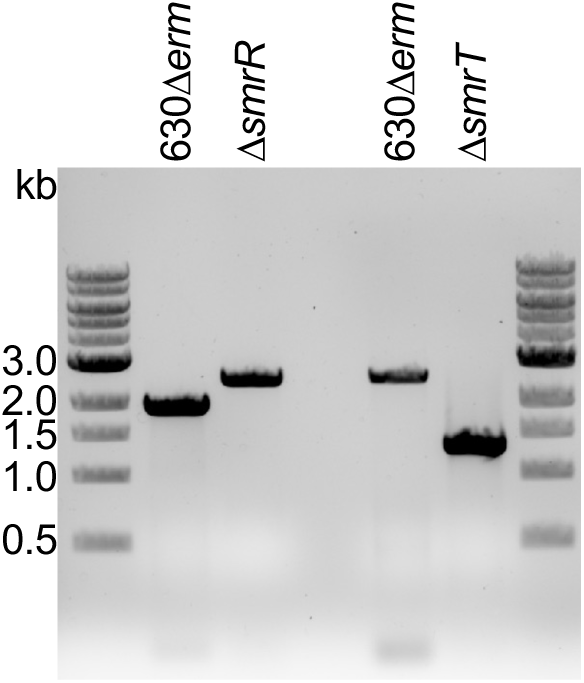
Verification of &*smrR* and &*smrT* mutant strains. PCR products were generated using flanking primers for *smrR* (*CD2505,* TetR-like regulator), or *smrT* (*CD2506,* MFS transporter). Genomic DNA from strain MC1681 (&*CD2505::ermB*), MC1682 (&*CD2506*) or 630—*erm* (control, wild type) were used as template, as indicated. The wild-type PCR product for *CD2505* is 2017 bp (primers oMC2321/oMC2324), and 2649 bp for &*CD2505* replaced by the *ermB* cassette. For *CD2506*, the wild type product is 2961 bp (primers oMC2541/oMC2544), and 1316 bp for &*CD2506*.

**Figure S3.**
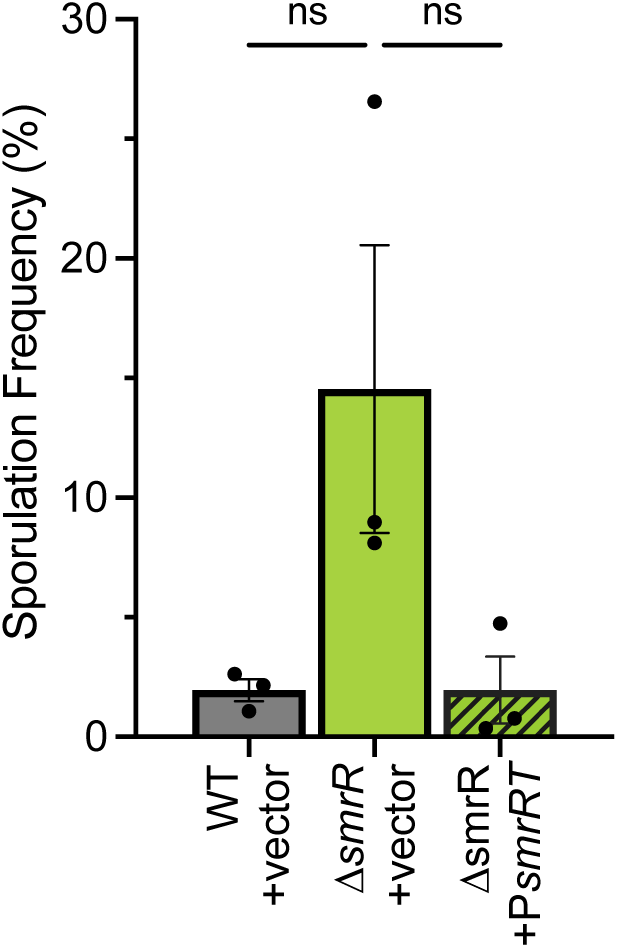
Complementation of the *smrR* mutant. Sporulation frequency of the wild-type control with empty vector (MC324; 630&*erm*, pMC123), the mutant with vector MC1771 (&*CD2505*, pMC123), and the mutant complemented with the *smr* operon expressed from the native promoter, MC1861 (&*CD2505*, pMC1059). Strains were grown in 70:30 broth, pH 7.2 with 2 *µ*g/ml thiamphenicol sporulation was determined after 24 h. The means, individual data points, and standard deviations are shown for three independent replicates. A one-way ANOVA with Sidak9s multiple comparisons test was used to compare the mutant to other strains. ns=not statistically significant

**Figure S4.**
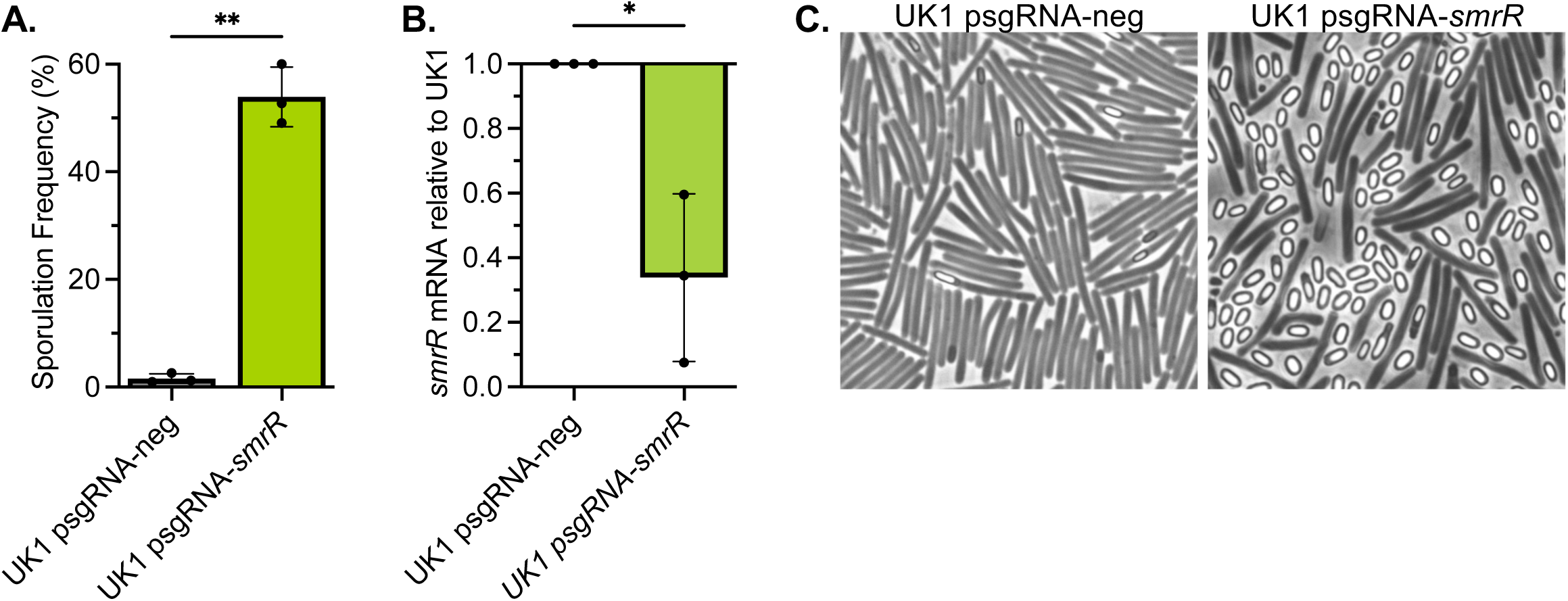
CRISPRi knockdown of *smrR* in strain UK1 increases sporulation and *smrT* expression. **A**) Ethanol-resistant spore formation of strain UK1 carrying a negative control vector p*sgRNA*-neg (MC2186) or p*sgRNA-smrR* (MC2263) grown 24 h on 70:30 agar with 2 *µ*g/ml thiamphenicol and 1 *µ*g/ml nisin. **B)** qRT-PCR analysis of *smrR (CDR20291_2397)* expression in MC2186 and MC2263 on 70:30 sporulation agar (H_12_). **C)** Phase-contrast micrographs of representative samples from the sporulation assay shown in A. The means, individual data, and standard deviations for three biological replicates are shown. \**P* f 0.05, \*\**P* f 0.01 by Student9s *t-*test.

**Figure S5.**
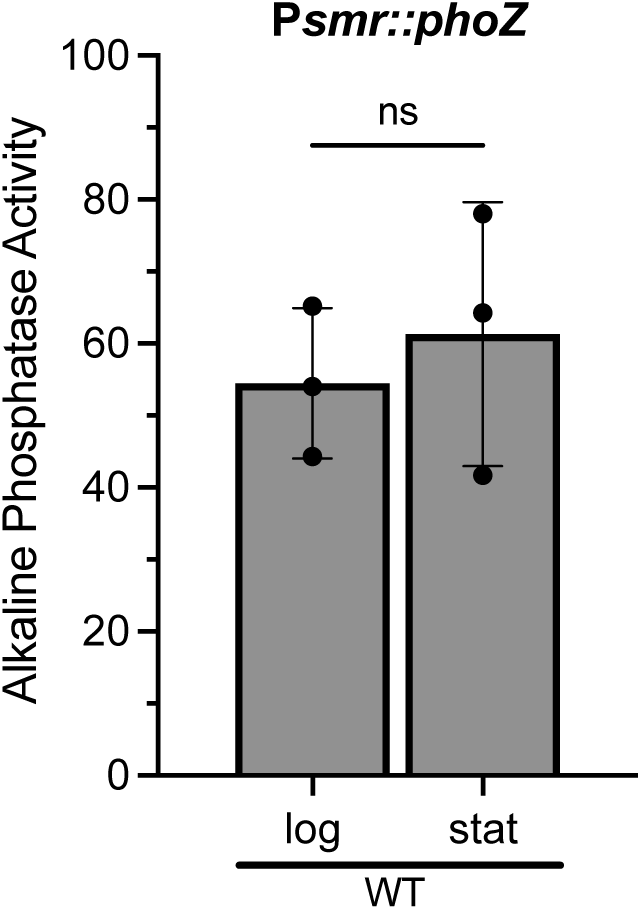
Expression from the *smr* promoter is independent of growth stage. Alkaline phosphatase activity of the P*smr::phoZ* fusion in *C. difficile* strain 630&*erm* (WT, MC1775). Strain was grown in 70:30 broth, pH 6.2 with 1 *µ*g/ml thiamphenicol and samples were taken during logarithmic growth (log, OD_600_ 0.5) and early stationary phase (stat, OD_600_ 1.0). The means, individual data points, and standard deviations are shown for three independent replicates. Data were analyzed by paired Student9s *t* test. \**P* f 0.05; ns: not significant.

**Figure S6.**
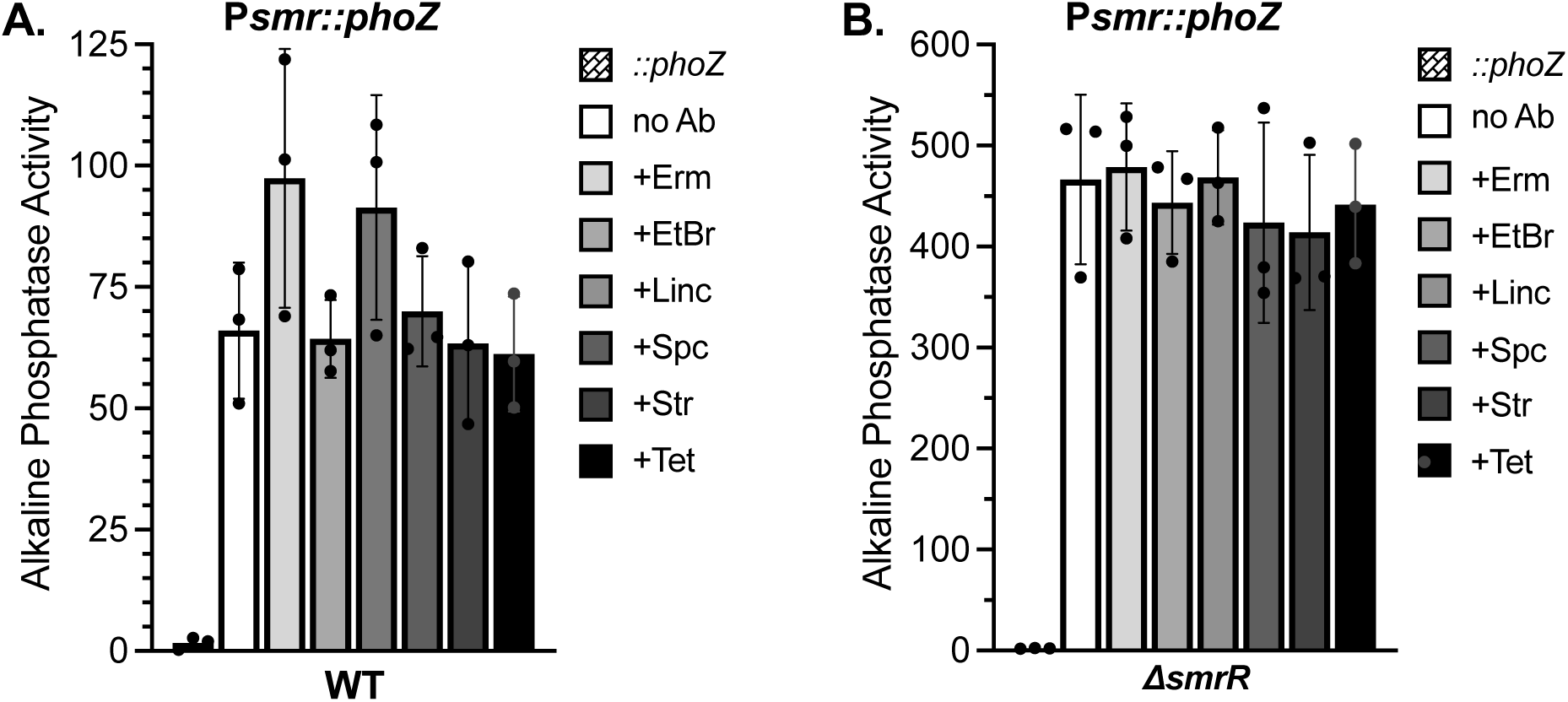
Erythromycin and lincomycin modestly induces expression from the *smr* promoter through SmrR. Alkaline phosphatase activity of the P*smr::phoZ* fusion in **A)** *C. difficile* strain 630&*erm* (WT, MC1775) and a promoterless control (*::phoZ,* MC448) or **B)** the &*smrR* mutant (MC1777) and a promoterless control (*::phoZ,* MC1776). Strains were grown in 70:30 broth, pH 6.2 with 1 *µ*g/ml thiamphenicol, with or without Erm (0.1 *µ*g/ml erythromycin), EtBr (0.5 *µ*g/ml ethidium bromide), Linc (0.5 *µ*g/ml lincomycin), Spc (10 *µ*g/ml spectinomycin), Str (10 *µ*g/ml streptomycin), or Tet (0.1 *µ*g/ml tetracycline). Samples were taken during logarithmic growth (OD_600_ 0.5). The means, individual data points, and standard deviations are shown for three independent replicates. Data were analyzed by one-way ANOVA comparing +antibiotic to the no antibiotic control. ns: not significant.

**Supplemental Table S1.**
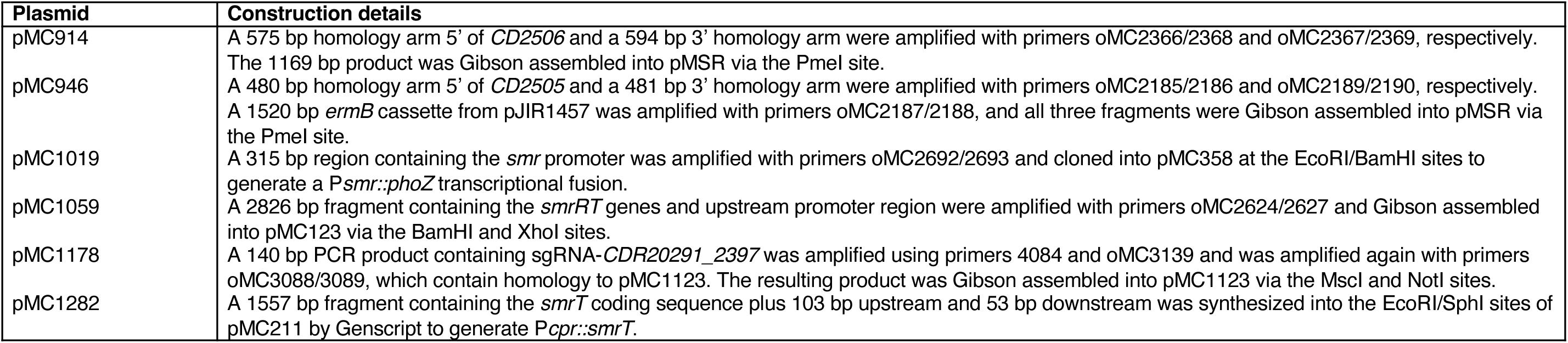
Vector construction.

